# SynBOLD-DisCo: Synthetic BOLD images for distortion correction of fMRI without additional calibration scans

**DOI:** 10.1101/2022.09.13.507794

**Authors:** Tian Yu, Leon Y. Cai, Victoria L. Morgan, Sarah E. Goodale, Dario J. Englot, Catherine E. Chang, Bennett A. Landman, Kurt G. Schilling

**Affiliations:** Department of Computer Science, Vanderbilt University, Nashville, TN, USA; Department of Biomedical Engineering, Vanderbilt University, Nashville, TN, USA; Department of Radiology and Radiological Sciences, Vanderbilt University Medical Center, Nashville, TN, USA; Vanderbilt University Institute of Imaging Science, Vanderbilt University Medical Center, Nashville, TN, USA; Department of Electrical and Computer Engineering, Vanderbilt University, Nashville, TN, USA; Department of Neurological Surgery, Vanderbilt University Medical Center, Nashville, TN, USA

**Keywords:** functional MRI, blood oxygen level dependent (BOLD) signal, susceptibility-induced distortion correction, image synthesis, 3D U-net, topup, Synb0-DisCo

## Abstract

The blood oxygen level dependent (BOLD) signal from functional magnetic resonance imaging (fMRI) is a noninvasive technique that has been widely used in research to study brain function. However, fMRI suffers from susceptibility-induced off resonance fields which may cause geometric distortions and mismatches with anatomical images. State-of-the-art correction methods require acquiring reverse phase encoded images or additional field maps to enable distortion correction. However, not all imaging protocols include these additional scans and thus cannot take advantage of these susceptibility correction capabilities. As such, in this study we aim to enable state-of-the-art distortion correction with FSL’s *topup* algorithm of historical and/or limited fMRI data that include only a structural image and single phase encoded fMRI. To do this, we use 3D U-net models to synthesize *undistorted* fMRI BOLD contrast images from the structural image and use this undistorted synthetic image as an anatomical target for distortion correction with *topup*. We evaluate the efficacy of this approach, named SynBOLD-DisCo (synthetic BOLD images for distortion correction), and show that BOLD images corrected using our approach are geometrically more similar to structural images than the distorted BOLD data and are practically equivalent to state-of-the-art correction methods which require reverse phase encoded data. Future directions include additional validation studies, integration with other preprocessing operations, retraining with broader pathologies, and investigating the effects of spin echo versus gradient echo images for training and distortion correction. In summary, we demonstrate SynBOLD-DisCo corrects distortion of fMRI when reverse phase encoding scans or field maps are not available.

## 1. INTRODUCTION

The blood oxygen dependent (BOLD) signal in functional magnetic resonance imaging (fMRI) is an established way of assessing neuronal activity in the brain by measuring the coupling of such activity with associated hemodynamic responses [1]. By evaluating changes in blood flow in different parts of the brain, fMRI BOLD images allow researchers to investigate associations of localized brain activity with cognition, sensation, and overall health [1], [2]. Further, by evaluating how BOLD signals correlate across different regions of the brain, fMRI BOLD images provide investigators a picture of the functional connections in the brain, both at rest and in response to tasks [1], [3]. These advances have yielded significant discoveries in neuroscience, such as the identification of the default mode network and an improved understanding of brain changes due to neurological conditions, including Alzheimer’s disease and epilepsy [4]–[7].

One critical limitation of fMRI BOLD images arises due to the echo planar imaging (EPI) acquisition schemes used to capture them. These schemes suffer from increased sensitivity to susceptibility-induced off resonance fields and magnetic field inhomogeneity, particularly in the phase encoding direction [8]–[10]. The result of this is that fMRI BOLD images acquired with EPI acquisitions can contain geometric distortions that result in inaccurate localization of voxels and subsequently confound spatial hemodynamic findings in the brain [8].

Reducing or correcting EPI distortions generally requires sequence modification (i.e., parallel imaging, increased bandwidth, improved shimming protocols) or post-acquisition image processing [8], [11]–[17]. While there are several strategies to correct distortions in image processing pipelines, including using b0 field maps or nonlinear registration to structural images, one technique that has risen to prominence in recent years is the use of reverse phase encoded images (i.e. using two scans with opposite phase encoding direction and hence equal and opposite distortions) in order to estimate and correct distortions [16]. For example, to correct distortions in an image phase encoded in the anterior-to-posterior (AP) direction, a reverse phase encoded image in the posterior-to-anterior (PA) direction is required. This type of correction is implemented in various software packages, (i.e., the *topup* algorithm in the FSL toolkit) and has several advantages over field maps and nonlinear registrations, including direct empirical estimates of distortion and appropriate signal intensity corrections [16], [17]. Additionally, correction with reverse phase encoded images, though typically recommended for spin echo images [18], may also be performed on gradient echo BOLD images directly where susceptibility fields can cause both distortion and unrecoverable signal loss [19].

One key limitation of this approach is that reverse phase encoded images are not always acquired due to time constraints, acquisition difficulties or artifacts, or unawareness of these processing techniques. Additionally, these images were not always acquired in historical datasets. Thus, there is a gap in the field where a method for correcting susceptibility-induced geometric distortions without these additional calibration scans is needed.

In this study, we aim to enable *topup*-like processing with historical and/or limited fMRI data that includes only the distorted forward phase encoded functional data and an associated structural image. Inspired by image synthesis in both diffusion [20], [21] and functional MRI literature [22], we leverage deep learning to synthesize an *undistorted* BOLD image from the acquired *distorted* BOLD image and a T1 weighted (T1w) structural scan. This undistorted, synthetic image with BOLD contrast can be used as a calibration scan to perform distortion correction of a distorted BOLD image by inputting both into *topup* and informing the algorithm that the synthesized image has an infinite bandwidth in the phase encoding direction (i.e., it is undistorted). We name this pipeline SynBOLD-DisCo (synthetic BOLD images for distortion correction). We show that this pipeline corrects image geometry in regions most susceptible to distortions, results in better matching to the structural image, and is practically equivalent to running *topup* with both forward and reverse phase encoded data.

## 2. METHODS

Figure 1 depicts our proposed pipeline based on our prior related work in diffusion MRI, Synb0-DisCo [20], [21]. The objective is to use an averaged distorted BOLD signal and a T1w anatomical MRI to generate a synthetic undistorted BOLD signal with appropriate geometry and contrast. This synthetic volume can then serve the role of a reverse phase encoded scan or field map with infinite bandwidth, providing state-of-the-art correction methods (i.e., FSL’s *topup*) the necessary information to correct the distorted image [16], [17].

**Figure 1.**
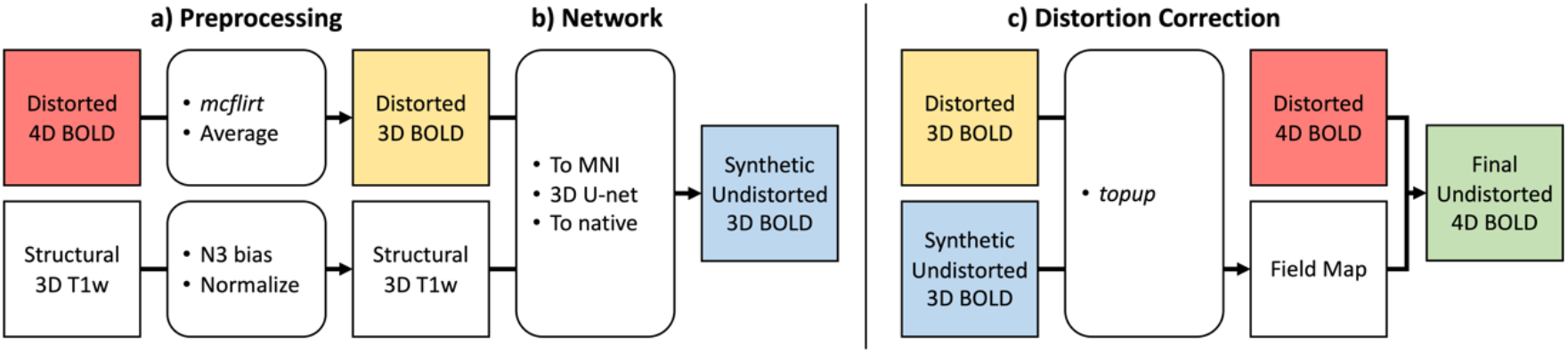
The SynBOLD-DisCo pipeline. (a) A distorted fMRI BOLD series and a T1w MRI are preprocessed and transformed to atlas space, (b) where a 3D U-net synthesizes a synthetic, undistorted fMRI BOLD image. (c) This synthetic image can be used with existing state-of-the-art distortion correction methods, like FSL’s *topup*, to compute field maps that allow the entire fMRI BOLD sequence to be corrected.

### 2.1 Data

Our aim is to train our network using a range of datasets, acquired on different scanners, with varying resolutions, contrasts, and distortion magnitudes. To this end, we utilize de-identified data from 9 different datasets (Table 1), each with a varying number of subjects, sessions, and complementary phase encoded image pairs (i.e., one forward and one reverse phase encoded scan) per session. This results in a total of 265 subjects and 805 fMRI BOLD image pairs that we subsequently use for training, validation, and testing.

**Table 1.** Datasets used for training, validating, and testing SynBOLD-DisCo.

| Dataset | Version | Subjects | Sessions | Images | Vendor | TE/TR (ms) |  | Resolution (mm) |  |
| --- | --- | --- | --- | --- | --- | --- | --- | --- | --- |
|  |  |  |  |  |  | T1w MRI | fMRI | T1w MRI | fMRI |
| ds002603 | 1.0.0 | 23 | 1 | 92 | Philips | 3.70/8.10 | 25-35/550-2000 | 1 iso | 2-3 iso |
| ds002606 | 1.1.0 | 10 | 2 | 20 | Siemens | 3.40/2.60 | 22/1228 | 0.8 iso | 1.6 iso |
| ds002655 | 1.0.1 | 35 | 1 | 35 | Siemens | 2.26/1900 | 45.2/2200 | 0.97x0.97x1 | 1.8 iso |
| ds002685 | 1.3.1 | 12 | 2-14 | 363 | Siemens | 3.16/2300 | 27/2000 | 1 iso | 1.5 iso |
| ds002738 | 1.0.2 | 33 | 1 | 33 | GE | —/— | 30/680 | 0.9 iso | 2.2 iso |
| ds002878 | 2.0.0 | 35 | 1 | 35 | Siemens | 2.10/— | 30/1400 | 0.875 iso | 2.5 iso |
| ds003037 | 1.0.1 | 53 | 1-6 | 138 | Philips | 3.50/7.00 | 30/2000 | 1 iso | 3x3x3.33 |
| ds003752 | 1.0.0 | 3 | 1 | 15 | Siemens | 2.20/2.40 | 37/800 | 0.8 iso | 2 iso |
| VUIIS | — | 61 | 1-2 | 74 | Philips | 3.00/8.00 | 23/2000 | 1 iso | 2.5 iso |

iso = isotropic, — = not applicable or not given.

Eight of the datasets are open-source and downloaded directly from OpenNeuro.org, while one dataset was acquired at the Vanderbilt University Institute of Imaging Science (VUIIS) under Institutional Review Board (IRB) 182089 and 181231. Importantly, these datasets were selected based on an exhaustive review of datasets available on OpenNeuro.org that contain the necessary complementary phase encoded BOLD data to train our model against gold standard distortion correction. The digital object identifier (DOI) for OpenNeuro.org dataset ds123456 version 7.8.9 follows doi:10.18112/openneuro.ds123456.v7.8.9.

In addition, to externally validate SynBOLD-DisCo on a dataset with no constituent data included in training, we leverage a cohort acquired at the National Institutes of Health under protocol 00-N-0082 and analyzed at VUIIS under IRB 181540. These data, referred to as the “external validation” cohort, consist of 13 subjects with T1w MRI and fMRI BOLD images (either resting state, task-based, or both) without reverse phase encoded scans. Thus, these data cannot undergo traditional distortion correction and serve as an initial external validation of the efficacy of the proposed pipeline. The T1w MRI were acquired with a TE/TR of 4.25/2200ms at 1mm isotropic resolution and the fMRI were acquired with a TE/TR of 29.4/2100ms at 3×3×4mm or 3.2×3.2×4mm resolution.

### 2.2 Preprocessing

To prepare for training, we preprocess each complementary phase encoded BOLD image pair and associated T1w MRI to obtain a normalized T1w MRI and distorted mean BOLD signals as well as undistorted mean BOLD signals estimated by *topup* to act as training targets. These images are all registered and transformed to Montreal Neurological Institute (MNI) atlas space [23].

First, each image in the complementary phase encoded pair is motion corrected and averaged with FSL’s *mcflirt*, generating distorted complementary mean BOLD signals [24]. Second, we utilize the state-of-the-art distortion correction tool, *topup*, to take these complementary mean BOLD signals and generate a distortion correction field map [16], [17]. Undistorted mean BOLD signals (i.e., the training targets) are obtained by applying the correction field maps to the original complementary phase encoded images and taking the average volume, producing two corrected images from each complementary BOLD image pair.

Next, N3 bias field correction and intensity normalization are performed on the input T1w MRI image using the FreeSurfer library [25], [26]. To ensure the network inputs (i.e., the distorted mean BOLD image and normalized T1w MRI) and target (i.e., the undistorted mean BOLD image) are in the same space, FSL’s *epi_reg*, a rigid-body 6 degree of freedom transformation, is used to register both BOLD images to the T1w MRI [17]. The T1w MRI is then affine registered using the ANTS library to the 1.0 mm isotropic MNI ICBM 152 asymmetric template [27]. Subsequently, all images are transformed to MNI space at 2.5 mm isotropic resolution.

To prepare for training, the intensities of the normalized T1w MRI and the undistorted and distorted mean BOLD signals are scaled. We map the Freesurfer normalized T1w MRI intensities between 0 and 150 to between −1 and 1. An intensity of 0 and the 99^th^ percentile intensity of the distorted and undistorted mean BOLD signals are mapped to between −1 and 1.

Of note, the preprocessing pipeline at deployment differs slightly: undistorted mean BOLD signals estimated by *topup* from complementary phase encoded image pairs (i.e., the training targets) are not needed. However, the T1w MRI and distorted BOLD series undergo the same preprocessing described herein (Figure 1).

### 2.3 Network architecture, training protocol, and loss definition

Following Synb0-DisCo, a 3D U-net is used to generate a synthetic, undistorted image with BOLD contrast in 2.5 mm atlas space from the distorted mean BOLD signal and the T1w MRI, leveraging the *topup* undistorted mean BOLD images as targets [20], [21]. In brief, the network is based on the original U-net implementation with a dual channel input, leaky ReLU activations, and instance normalization instead of batch normalization layers.

The data are partitioned by subject for the test, validation, and training sets. We randomly withhold 12% of subjects for each dataset, resulting in 30 subjects (81 image pairs) and utilize the remaining 235 subjects (724 image pairs) in 5-fold cross validation for training and validation. Of note, since there are only 3 subjects in ds003752, this dataset is omitted from the testing set.

Each fold is trained for 120 epochs with a learning rate of 0.0001. Beta values of 0.9 and 0.999 are used with the Adam optimizer. A weight decay rate of 0.00001 is applied. After each epoch, the validation loss is computed, and the weights of the model are saved if it improves. This results in five trained networks, one for each fold. NVIDIA GeForce GTX 1080 Ti and Quadro RTX 5000 GPUs are used for training, requiring about 9GB of memory.

During each iteration, the following loss computation occurs. First, both the forward and reverse distorted mean BOLD signals (*x*_*f*_and *x*_*r*_, respectively) are passed through the network with the associated normalized T1w MRI to produce two synthetic, undistorted images with BOLD contrast (*y*_*f*_, and *y*_*f*_, respectively). Next, we compute three losses, denoting the forward and reverse state-of-the-art undistorted mean BOLD signals (i.e., the targets) as *z*_*f*_and *z*_*r*_, respectively: *MSE*(*y*_*f*_, *z*_*f*_), *MSE*(*y*_*r*_, *z*_*r*_), and *MSE*(*y*_*f*_, *y*_*r*_), where *MSE* is the mean squared error across the image voxels. We sum these three terms to produce the final loss. The rationale behind this is to enforce (1) that the synthesized images be similar to their respective undistorted targets and (2) that the forward and reverse images produce similar synthetic scans, similar to Siamese and null space network designs [20], [21].

### 2.4 Distortion correction leveraging the synthetic volume

All five networks (one from each fold) are used to make an inference. The final synthetic BOLD image is determined by the ensemble average of the predictions of the five networks. Inverse transformation is then applied to move the synthetic BOLD image back to the original BOLD space. The synthetic BOLD image is then passed into *topup* along with the forward or reverse mean distorted BOLD signal to generate a distortion correction field map which is subsequently used to correct the entire distorted BOLD signal. This results in the final output of the pipeline.

Notably, the synthetic image is considered to have infinite bandwidth (readout time of 0) for *topup* while the distorted can have arbitrary bandwidth relatively. Last, in order to match the smoothness of the synthetic image, the distorted image is smoothed slightly with a Gaussian kernel with standard deviation 1.15 mm prior to *topup* [20], [21]. The smoothness of the synthetic image originates from the network operating at a slightly lower resolution (2.5 mm isotropic) due to GPU memory constraints and interpolation while resampling back to subject space.

### 2.5 Quantitative evaluation of pipeline

To evaluate the effectiveness of our pipeline, metrics such as mean absolute percent difference and mutual information are used. For all subjects in the testing set, we calculate the mean absolute percent difference by comparing the distorted mean BOLD signals (“Distorted”), corresponding synthetic BOLD signals (“Synthetic”), and resulting undistorted mean BOLD signals from the pipeline (“SynBOLD”) with state-of-the-art *topup* correction with complementary phase encoded images (“Topup”) to assess both distortion correction accuracy and contrast accuracy. Second, mutual information is calculated by comparing the “Distorted”, “Topup”, “Synthetic”, and “SynBOLD” images with N3 bias corrected T1w weighted MRI to assess geometric similarity. For both experiments, all images are rigidly registered to T1w MRI image space and all metrics are computed within a brain mask. For the “SynBOLD” images, the mean undistorted BOLD signal is used to maintain comparisons in 3 dimensions. Pair-wise Wilcoxon sign rank tests with Bonferroni correction at 0.05 significance are used to determine statistical significance.

## 3 RESULTS

### 3.1 Network convergence and loss curves

We determine convergence of the dual channel U-net across all 5 folds through analysis of the loss curves (Figure 2). We observe a deep descent of the training loss within the first 30 folds and small improvements afterwards. Looking at the final 30 folds, we observe continued descent of the training loss with no improvement in the validation loss, suggesting convergence. The testing loss at the end of training was in line with the validation loss, suggesting appropriate generalizability.

**Figure 2.**
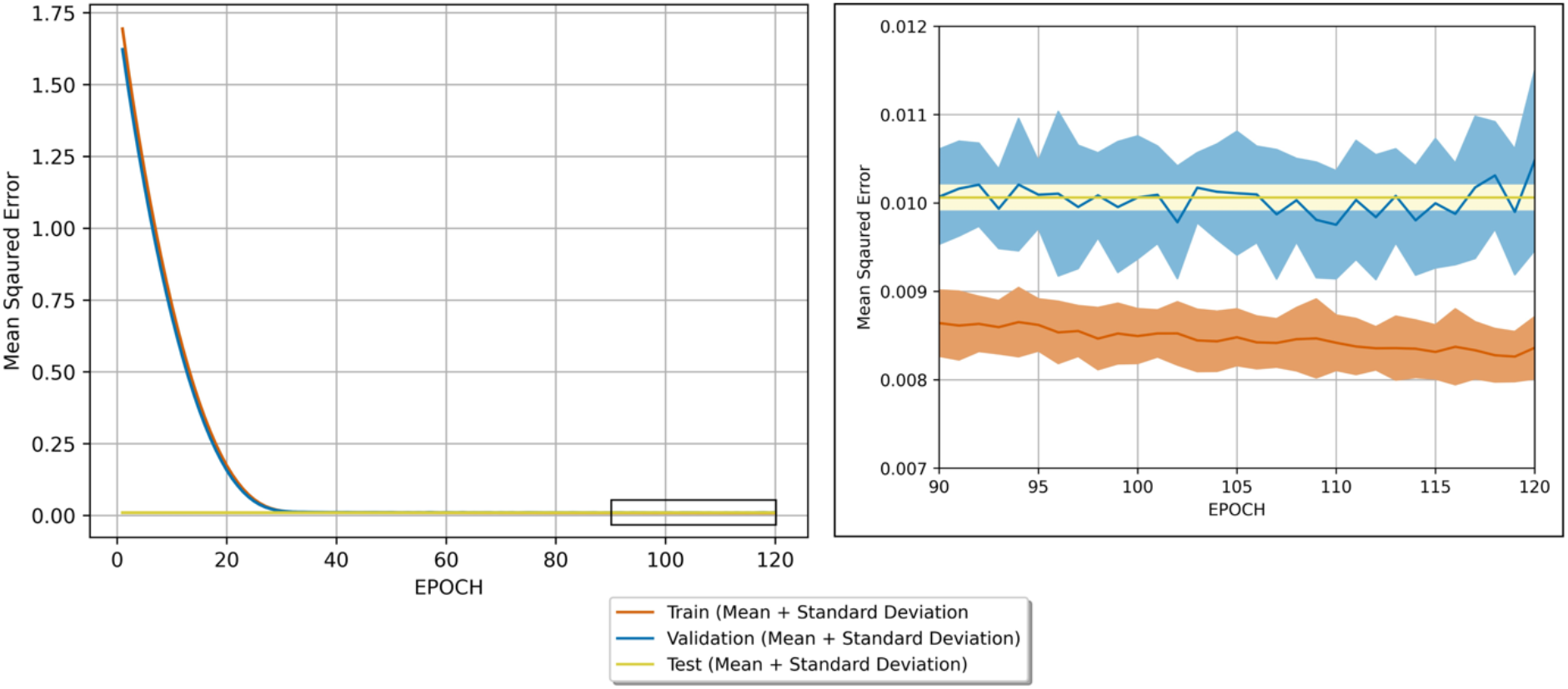
Training, validation, and withheld testing set losses. The mean and standard deviation of the losses for each dataset across the 5-folds are plotted and suggest network convergence and appropriate generalizability to withheld data by the end of training (120 epochs). Each of the five folds fulfilled early stopping criteria and the final models were saved between the 90^th^ and 120^th^ folds. The testing loss was computed once at the end of training.

Additionally, the plotted loss is a combination of three *MSE* losses, and the square root of the observed loss for the testing and validation sets hovers around 0.1. As the target images were normalized to between −1 and 1, we note the overall validation and testing error on the image level falls around of 5% or less of the image range, suggesting appropriate agreement.

### 3.2 Quantitative performance

We plot the mean absolute percent difference of the “Distorted”, “Synthetic (Intermediate)”, and “SynBOLD (Final)” BOLD signals in the withheld test set against the corresponding “Topup (State-of-the-art)” volumes to determine contrast and intensity accuracy after correction with our pipeline (Figure 3a). We find both the intermediate synthetic volume generated by the neural network and the final corrected mean BOLD signal leveraging the synthetic volume obtain statistically significant smaller differences against the state-of-the-art but not between each other, suggesting that both the synthetic network output and final undistorted fMRI data from our pipeline match the state-of-the-art distortion correction generated from complementary phase encoded images.

**Figure 3.**
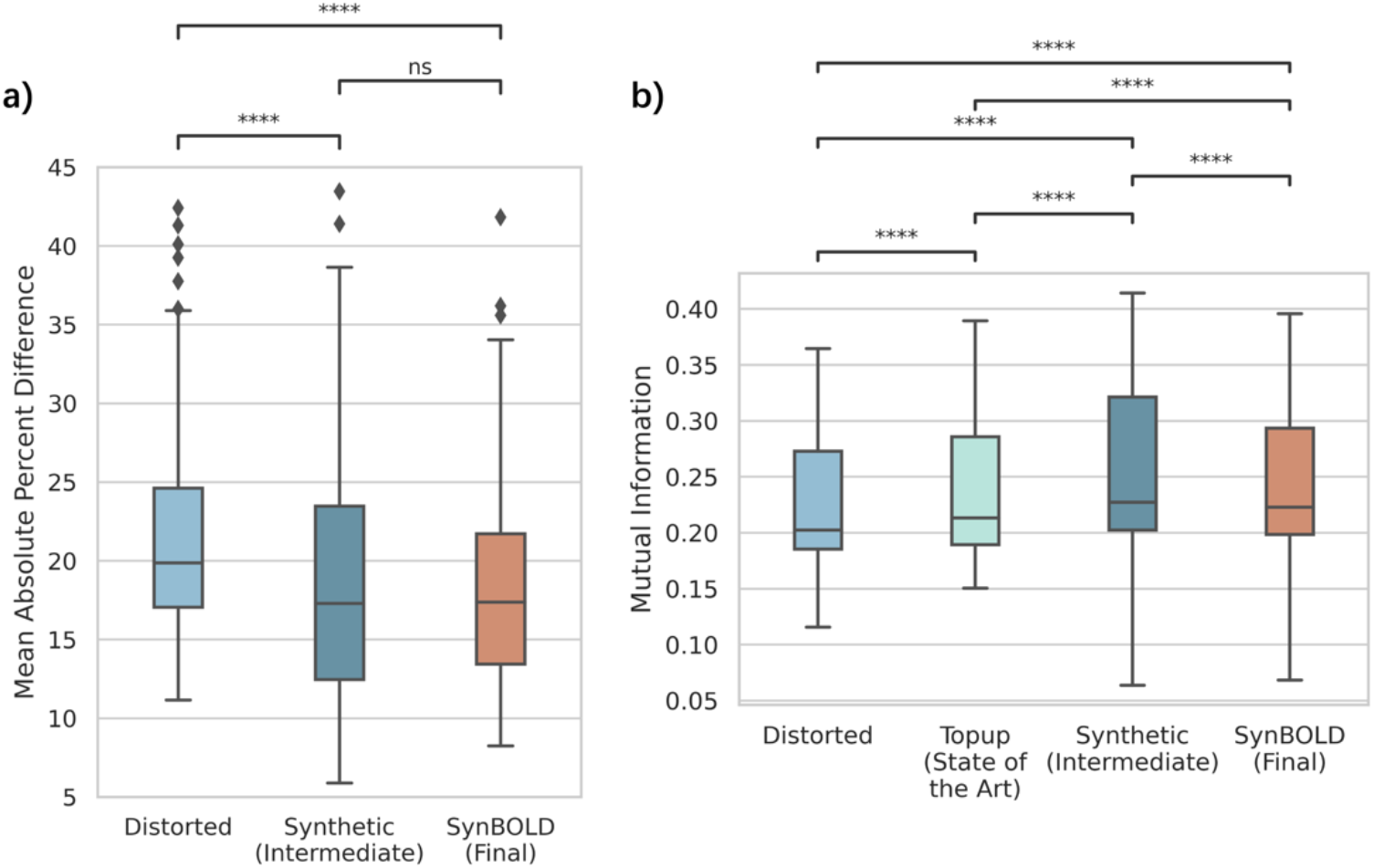
Quantitative evaluations of pipeline accuracy. (a) Mean percent difference between the “Distorted”, “Synthetic (Intermediate)”, and “SynBOLD (Final)” volumes used in our pipeline and the “Topup (State-of-the-art)” volume. Lower mean percent difference indicates increased similarity with the state-of-the-art. (b) Mutual information between the “Distorted”, “Topup (State-of-the-art)”, “Synthetic (Intermediate)”, and “SynBOLD (Final)” volumes and a corresponding N3 bias corrected T1w MRI. Higher mutual information indicates increased geometric similarity with the structural image. **** *p* < 0.0001 (Wilcoxon sign rank test with Bonferroni correction), ns = not significant.

To assess geometric similarity to structural scans, we plot the mutual information within the brain between the N3 bias corrected T1w MRI and each of the BOLD images used in the pipeline, “Distorted”, “Topup (State-of-the-art)”, “Synthetic (Intermediate)”, and “SynBOLD (Final)”, for each sample in the withheld test set to assess geometric accuracy (Figure 3b). We find the synthetic volume has the highest mutual information, followed by the SynBOLD-DisCo output, the state-of-the-art, and finally the distorted volume.

### 3.3 Qualitative performance

To visually understand the effects of the pipeline against structural imaging and the state-of-the-art, we visualize a representative sample in the withheld test set (Figure 4). We observe visually appreciable and comparable improvements in distortion correction with both our pipeline and the state-of-the-art as well as increased similarity to structural T1w MRI. We also confirm structural similarity of the synthetic intermediate volume produced by the 3D U-net with the T1w MRI.

**Figure 4.**
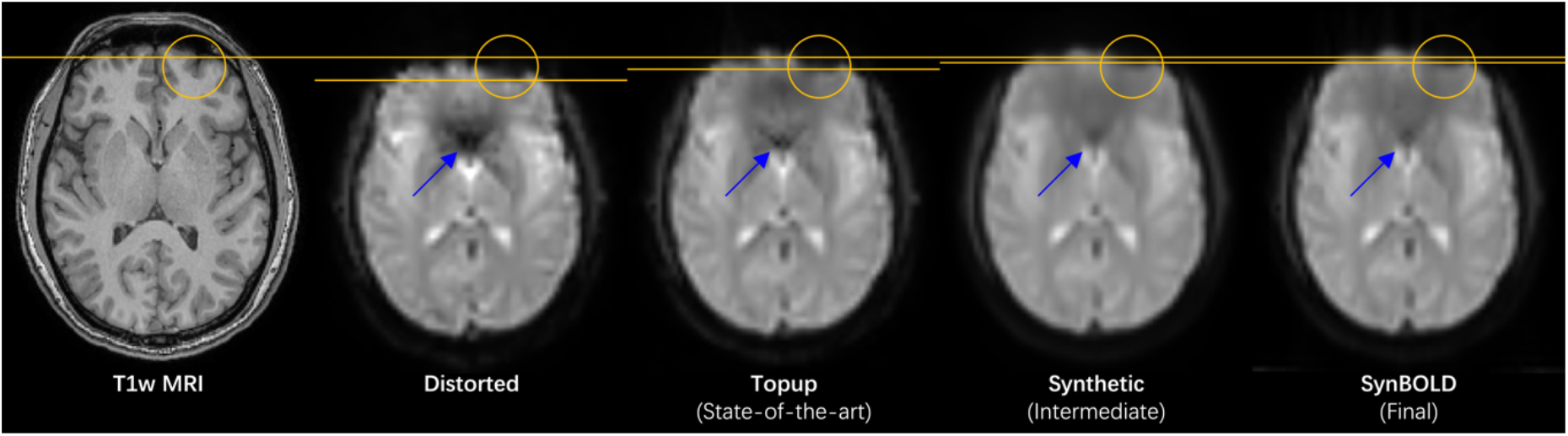
Visualization of a representative sample in the withheld test set, including the “Distorted”, “Topup (State-of-the-art)”, “Synthetic (Intermediate)”, and “SynBOLD (Final)” volumes and a corresponding structural T1w MRI. The yellow lines denote the anterior borders of the corresponding distorted sulci highlighted by the yellow circles. The blue arrows denote geometrically similar areas with improved signal intensities with distortion correction.

Additionally, we provide representative examples from each of the 8 datasets included in the withheld cohort in Figure 5. Importantly, our approach visually generalizes well across independent datasets, with results similar to state-of-the-art processing and geometry matching the anatomical images.

**Figure 5.**
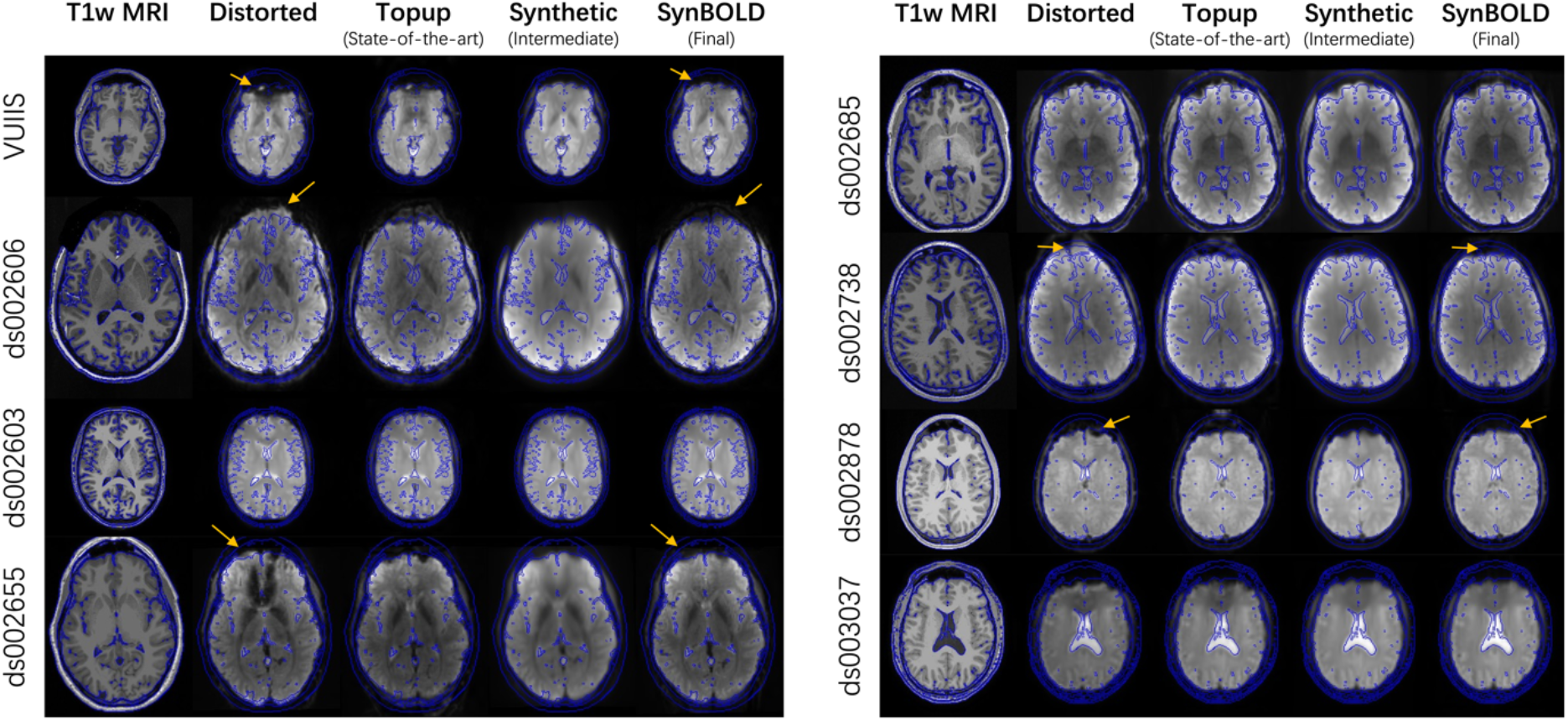
Visualization of a representative sample from each of the 8 withheld datasets, including the “Distorted”, “Topup (State-of-the-art)”, “Synthetic (Intermediate)”, and “SynBOLD (Final)” volumes and a corresponding structural T1w MRI. Blue lines indicate tissue interfaces computed on the T1w MRI. Yellow arrows denote examples of geometrically improved distortions with our pipeline.

### 3.4 External validation

We apply our pipeline to the external validation cohort acquired without phase encoded scans to evaluate its ability to correct images that are unable to undergo traditional state-of-the-art distortion correction with *topup*. We find in a representative sample that our pipeline produces images that are more geometrically similar to anatomical T1w MRI than the distorted input (Figure 6).

**Figure 6.**
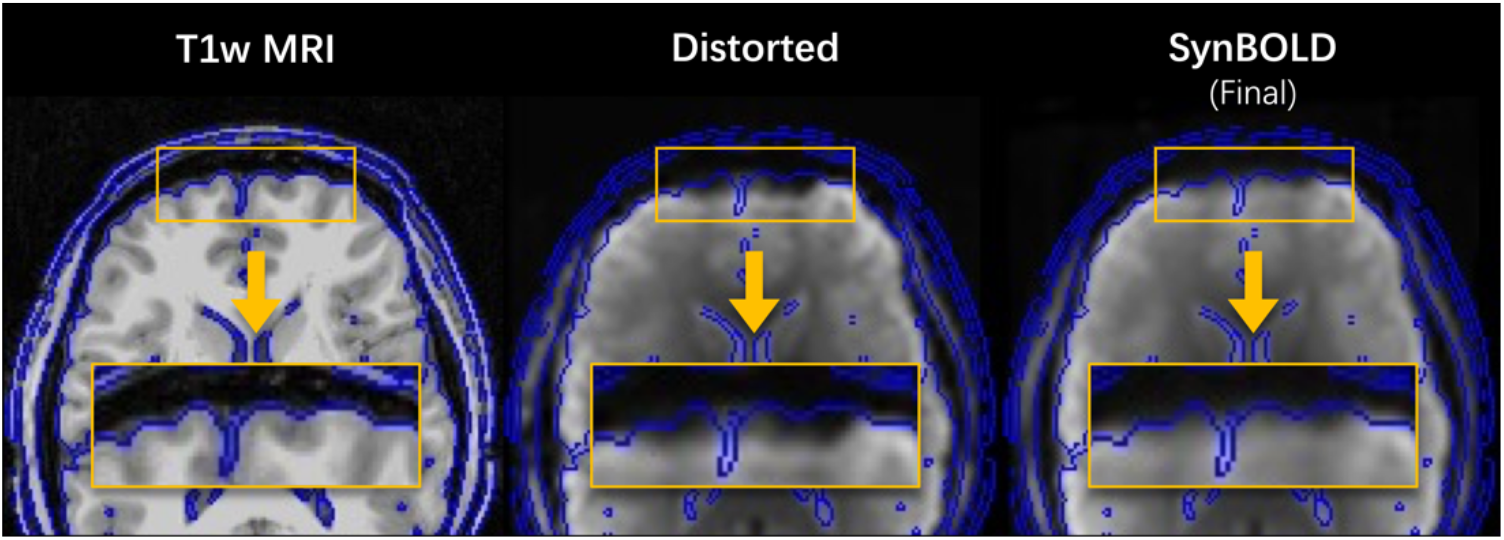
Visualization of a representative sample in the external validation set, including the “Distorted” input and “SynBOLD (Final)” output of our pipeline as well as a corresponding structural T1w MRI. Blue lines indicate tissue interfaces computed on the T1w MRI. The yellow boxes denote corresponding frontal gyri with improved geometric similarity to T1w MRI after SynBOLD. The “Topup (State-of-the-art)” volume is omitted because these data were acquired without reverse phase encoded images and thus are unable to undergo traditional distortion correction.

## 4. DISCUSSION

In this work, we demonstrate that distorted fMRI BOLD images and T1w MRI can be used to learn synthetic, undistorted volumes that facilitate susceptibility-induced distortion correction using established state-of-the-art methods without the need for reverse phase encoded scans or additional field maps. We determine this qualitatively by visualizing the distorted inputs, the synthetic volumes, and the pipeline outputs against both images corrected using reverse phase encoded scans and structural T1w MRI. These qualitative findings are supported quantitatively by reduced error between our pipeline outputs and the state-of-the-art as well as increased structural similarity between our pipeline outputs and T1w MRI. Importantly, we also demonstrate that our pipeline produces visually improved distortions in fMRI data unable to undergo traditional distortion correction techniques. Overall, this represents a distinct step forward toward improving the quality of fMRI BOLD images prior to subsequent analysis without the need for potentially complex or unreliable calibration scans.

This pipeline was inspired by recent advances in image synthesis for diffusion MRI distortion correction [20], [21], using the same network architecture described here to synthesize undistorted images for *topup*-like correction. These techniques have provided the ability to reliably correct diffusion MRI data [28] in cases where field maps and reverse phase encoded scans were unavailable [29]–[33], as is the case for legacy datasets or intra-operatively during surgery [34]. We imagine similar use cases for the current pipeline to investigate functional findings in areas prone to susceptibility-induced distortions.

Similar pipelines using synthesized images for fMRI do exist. For example, the recent study by Montez et al. uses T1- and T2-weighted contrast to synthesize a high resolution and high signal-to-noise ratio BOLD contrast image for use as a template for nonlinear warping for distortion correction [22]. Outputs from their pipeline perform similarly (or outperform) other approaches utilizing field mapping. In our study, rather than using the synthesized image as a reference for nonlinear warping, we have chosen to use the synthesized image for *topup*-like processing where the distortions are estimated from the data based on physical principles (distortion directions, magnitude, signal intensity variations). While nonlinear registration is certainly commonly performed, it may not be optimal as it attempts to match BOLD contrast (with both distortions and signal dropout) to structural images and may not appropriately address signal intensity variation (signal pileup or signal stretching) due to distortions.

Our technique will also face similar challenges in areas of signal dropout due to the use of gradient echo images in the training set. The use of gradient echo versus spin echo images to calculate the *topup* field maps may result in poor estimates in areas with little to no signal. However, no signal will be available to shift even if field maps are perfectly estimated [18]. For this reason, we train our network to estimate the corrected gradient echo images directly, which in theory should be images that are distortion corrected but suffer from the same signal dropout (allowing *topup* to disentangle dropout from distortion). Further, recent work suggests that using gradient echo images directly for distortion correction of other gradient echo images may be beneficial [19].

Another prominent limitation is the inclusion of data only phase encoded in the AP or PA directions and data without obvious lesions, such as tumors or plaques. This could potentially result in SynBOLD-DisCo being not applicable to data that do not fit these assumptions. Another limitation is that the assessments of distortion correction done herein were performed after rigid registration to T1w MRI space. We pursued this approach to minimize confounding of these metrics due to improper alignment between images. However, registration processes are imperfect and may introduce biases in evaluation. Last, despite statistically significant improvements in distortion with SynBOLD-DisCo, we note the distortion correction capacity of *topup* is imperfect, thus introducing a performance ceiling for SynBOLD-DisCo.

There are many additional directions this work could take to improve the research landscape. One is further external validation of distortion correction findings on independent, unrelated datasets. This could take the shape of direct geometric- and intensity-based assessments like those included presently, or an investigation of improvement in effect size or significance among studies of disease. Additionally, the integration of SynBOLD-DisCo into other fMRI preprocessing pipelines to address distortion correction as well as motion and normalization concerns would be beneficial for the field. As mentioned previously, retraining of our model with consideration for images phase encoded in different directions or on a population that include lesioned brains may be helpful in promoting SynBOLD-DisCo’s applicability to broader audiences. Last, investigating differences in spin echo versus gradient echo images for distortion correction would improve understanding of how these acquisitions impact fMRI BOLD analyses.

We make the SynBOLD-DisCo source code and trained weights available at github.com/MASILab/SynBOLD-DisCo along with containerized implementations to promote further investigation in the field.

## ACKNOWLEDGEMENTS

The authors thank Lillian Y. Cai for a thorough readthrough of the methodology. This work was conducted in part using the resources of the Advanced Computing Center for Research and Education at Vanderbilt University, Nashville, TN. This work was supported by the National Institutes of Health (NIH) under award numbers R01EB017230, RF1MH123201, R01NS110130, R01NS108445, R01NS112252, K01EB032989, T32EB001628, 1U34DK123895-01, U34DK123894-01, and T32GM007347 and by the National Science Foundation (NSF) under award number 2040462. This research was conducted with the support from the Intramural Research Program of the National Institute on Aging of the NIH. The content is solely the responsibility of the authors and does not necessarily represent the official views of the NIH or NSF.

